# Single-cell-resolved natural giant virus infections reveal conserved protist immunity

**DOI:** 10.64898/2026.10.11.758280

**Authors:** Amir Fromm, Tomas Alarcón-Schumacher, Christopher Bellas, Talia S. Shaler, Daniella Schatz, Max Emil Schön, Nele Schewe, Karel Kopejtka, Ruben Sommaruga, Frank O. Aylward, Matthias G. Fischer, Assaf Vardi

## Abstract

Single-cell eukaryotes (protists) are key players in microbial ecosystems. Yet, our understanding of how they interact with viruses in nature and how antiviral defenses shape these relationships remains limited. Using a single-cell multi-omics approach, we mapped associations between giant viruses and their hosts and identified active giant-virus infections in an alpine lake protist community. In naturally infected dinoflagellates, infection was accompanied by the selective induction of viperin, an ancient antiviral enzyme conserved from bacteria to humans. We also observed virus-encoded homologs of immunity-related genes whose expression correlated with viperin expression, raising the possibility of a role in viral counter-defense. These findings reveal an ecological dimension of protist immunity and provide a framework for tracing the host-virus arms race across divergent cellular lineages.

---

Viruses impose a major selective pressure on cellular life, driving the evolution of antiviral defense systems across bacteria and eukaryotes (*1–3*). Despite advances in our understanding of antiviral immunity in animals, plants, and bacteria (*1–3*), little is known about antiviral immune systems in protists, including their abundance and ecological importance.

This knowledge gap is especially relevant for protists infected by giant viruses. Giant viruses are widespread in aquatic environments and can strongly affect protist mortality, population dynamics, and evolution (*4*, *5*). Although environmental metagenomic sequencing has rapidly expanded the known diversity of giant viruses (*6–8*), the natural hosts of many viral lineages remain unknown, and little is known about the cellular defenses deployed against them in nature. Bulk metagenomics can reveal the diversity and biogeography of viral genomes, but generally cannot determine which host is infected, or distinguish active infection from extracellular viral DNA, ingested particles, or endogenous viral elements. As a result, viral infection dynamics and host responses are difficult to resolve in their native ecological context. Recent work has uncovered host responses associated with viral resistance in cultured algae (*9*). Nevertheless, endogenous antiviral defense systems against giant viruses remain underexplored, with known forms of host protection instead relying on third-party interactions, such as defensive symbiosis with bacteria (*10*) and the use of virophages (*11*). However, identifying antiviral defense systems or demonstrating their activity in model experimental systems still does not establish how hosts use them in their native environment. This ecological dimension is particularly inaccessible in uncultivated microbial eukaryotes, where both the infecting virus and host response must be resolved within the same cell.

Single-cell RNA sequencing (scRNA-seq) can bypass this limitation by linking the expression of viral and host genes in individually infected cells (*12*) and using transcriptomics to identify infection states (*13*, *14*). This enables the discovery of viral takeover strategies and host antiviral responses in natural populations, addressing a major gap in our understanding of both protist immunity and the evolutionary history of cellular defense. Furthermore, the vast genomic diversity of natural plankton virus interactions, far exceeding the narrow repertoire captured by laboratory model systems, provides an untapped resource for uncovering immune mechanisms and defining how they operate under ecologically relevant conditions.

Here, we combined single-cell RNA sequencing, single-cell DNA sequencing, and metagenomics to resolve giant-virus infections within a natural alpine lake protist community. Freshwater lakes in high altitudes are harsh environments whose viral diversity remains largely unknown. We identified diverse host-virus associations across various protist lineages, including eukaryotic taxa for which no viruses have been identified to date, as well as viral lineages with hitherto unknown hosts. We show that the evolutionarily conserved antiviral protein viperin, conserved from humans to bacteria, is expressed in single-celled eukaryotes in response to viral infection.

## Results and Discussion

### Multi-omics resolves protist community structure in an alpine lake

To resolve protist-giant virus interactions within their native ecological context, we sampled the microbial community of Gossenköllesee, a freshwater lake located at 2416 m elevation in the Austrian Alps, and combined single-cell RNA sequencing, single-cell DNA sequencing, and bulk metagenomics (Fig. 1; table S1). Bulk metagenomics provided a reference catalog of host genes and giant virus genomes, whereas single-cell approaches linked viral sequences to individual protist cells and distinguished viral association from active infection.

**Fig. 1.**
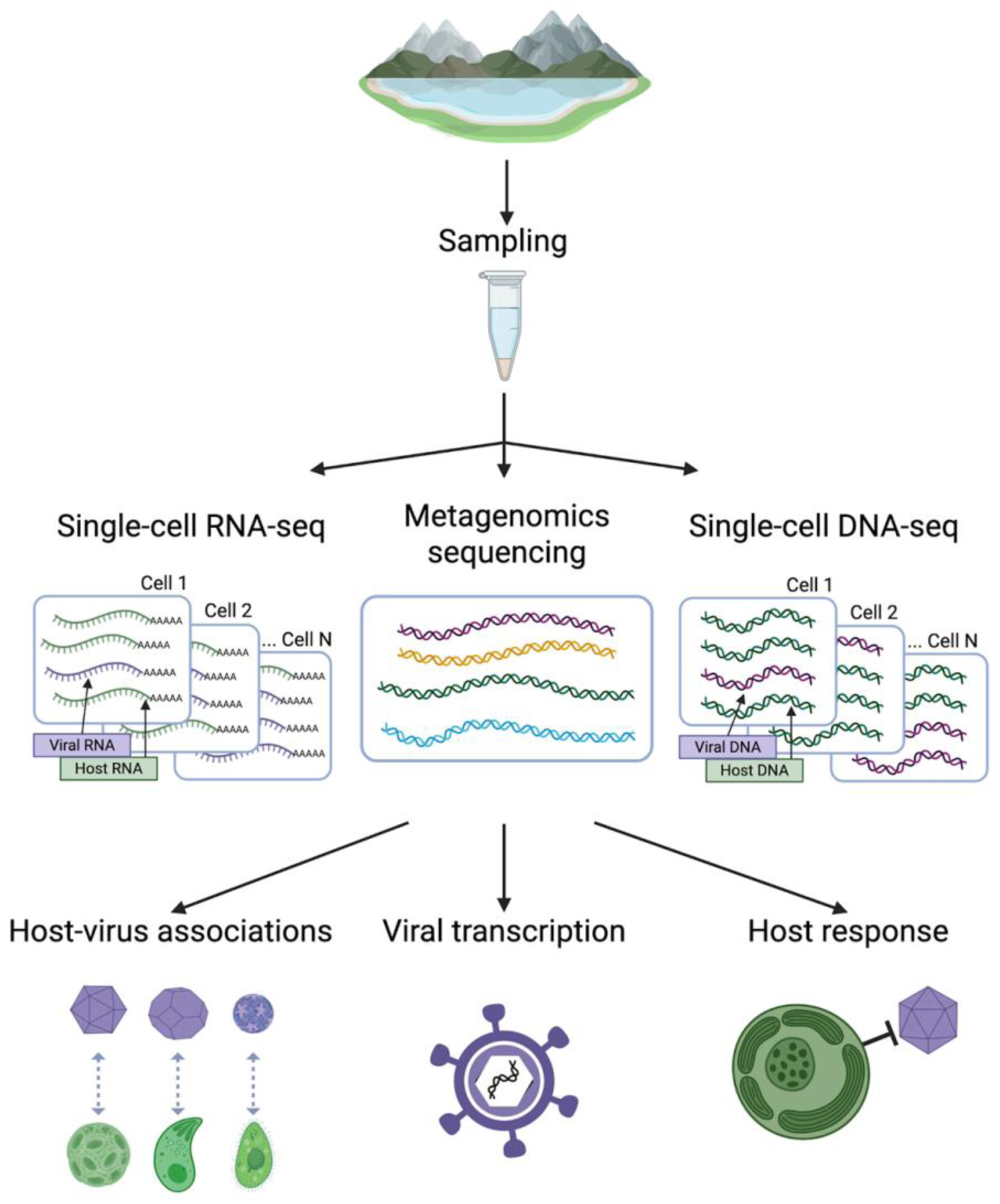
Multi-omics framework for studying giant virus-host interactions in natural microbial communities. Water samples from Gossenköllesee were divided for bulk metagenomic sequencing, single-cell DNA sequencing, and single-cell RNA sequencing. The figure was created with BioRender.com.

The complementary sequencing approaches provided a framework for linking viral diversity to individual host cells and their transcriptional states. Dinoflagellates (class Dinophyceae) dominated the eukaryotic fraction of the bulk metagenomes, representing 46% of reads mapped to 18S rRNA genes (Fig. 2A, data S1), followed by chrysophytes (11.5%), haptophytes (8%), and diatoms (class Bacillariophyceae) (7%).

**Fig. 2.**
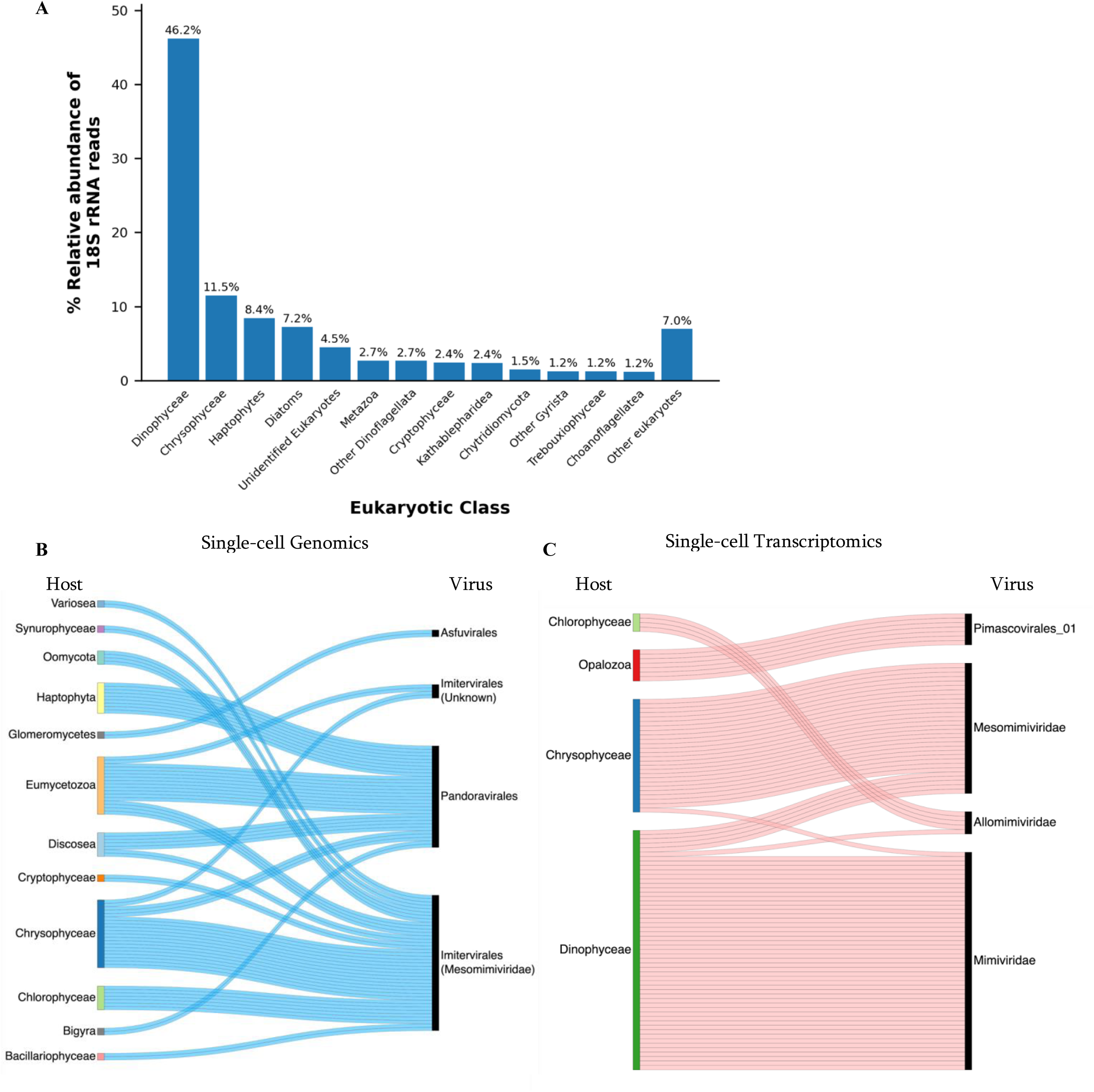
Eukaryotic community composition and single-cell pairing between giant viruses and their predicted protist hosts in Gossenköllesee. (A) Relative abundances of eukaryotic classes inferred from four bulk metagenomic samples. Relative abundance was calculated as the proportion of 18S rRNA reads assigned to each taxonomic group among all 18S rRNA reads classified as eukaryotic. (B-C) Single-cell pairing between giant viruses and their predicted protist hosts in Gossenköllesee. Connecting lines represent predicted host-virus associations identified in at least two single cells. (B) Predicted viruses infecting single host cells detected by scDNA-seq (n = 76). Associations were defined by the co-occurrence of host and virus DNA within individual cells after filtering for high-confidence viral assignments (see Methods). (C) Co-occurrence of virus and protist transcripts as detected by scRNA-seq. (n = 91).

### Single-cell genomics reveals diverse giant virus host associations

Single-cell DNA sequencing recovered 12,309 single amplified genomes (SAGs), of which 4,389 were classified as eukaryotic (data S2). Screening single-cell genomes for hallmark Nucleocytoviricota genes identified 650 SAGs containing ≥3 giant-virus marker genes such as major capsid protein and DNA polymerase (fig. S2; see Methods), thereby providing a substantial collection of draft viral genomes directly from individual cells. Importantly, their recovery does not depend on the availability of corresponding viral genomes assembled from bulk metagenomic data, highlighting the potential of scDNA-seq to independently capture viral genomic diversity while simultaneously providing virus-host associations at the single-cell level. We identified 76 protist cells that could be confidently associated with a single viral order (Fig. 2B, data S3). Predicted host-virus associations revealed a broad host range among the recovered giant virus lineages, though distinct taxonomic preferences were evident (Fig. 2B). Members of the order *Imitervirales*, primarily represented by the *Mesomimiviridae* (IM_01) family, were most frequently associated with Chrysophyceae (Fig. 2B, 15 associations), consistent with previous environmental observations (*15*) and the recent isolation of a chrysophyte-infecting imitervirad (*16*). Additional associations across diverse protist lineages suggest a broad host range for mesomimivirids in Gossenköllesee (Fig. 2B). In parallel, members of the proposed order “Pandoravirales” exhibited a diverse host range spanning multiple heterotrophic and photosynthetic lineages (Fig. 2B). Pandoravirads were associated with two amoebozoan lineages, Eumycetozoa (11 cells) and Discosea (5 cells), providing single-cell evidence for amoebae as natural hosts (Fig. 2B). Nine haptophyte cells also contained pandoravirad DNA, consistent with recent phylogenetic evidence demonstrating that the haptophyte-infecting *Coccolithoviridae* family is closely related to this lineage (*17*). The presence of viral DNA within a cell, however, does not necessarily indicate active infection and may also derive from ingested particles (i.e., “virovory”) (*18*) or endogenous viral elements (*19–22*). We therefore used single-cell transcriptomics to identify active infections in native cells.

### Single-cell transcriptomics reveals active giant virus infections in native protist hosts

Our scRNA-seq dataset comprised 7,127 cells with individual transcripts quantified using unique molecular identifiers (UMIs). Of these, 3,419 were taxonomically assigned using assembled 18S rRNA transcripts (data S4). To identify active viral infections, assembled transcripts from individual cells were queried against a custom database of giant-virus proteins derived from metagenome-assembled giant virus genomes (GVMAGs) assembled from the Gossenköllesee bulk metagenomes (see Methods). We identified 91 cells that were confidently assigned to a single viral family (Fig. 2C and data S5). In contrast to the broad associations detected by single-cell genomics, active viral transcription concentrated on four protist classes, each predominantly associated with a distinct giant virus lineage.

The most prevalent infection occurred in Dinophyceae, where 50 of 54 virus-positive cells were infected by mimivirids (Fig. 2C), consistent with previous observations of mimivirid infection during a dinoflagellate bloom (*23*). However, other studies have linked dinoflagellates to giant viruses from different lineages, including *Mesomimiviridae* (*24*) and *Asfarviridae* (*25*). Mesomimivirid infection was detected in 26 Chrysophyceae cells, consistent with scDNA-seq results (Fig. 2B), whereas four Chlorophyceae cells expressed allomimivirid genes, a group that was previously shown to infect green algae (*26*). Seven Opalozoa cells were infected by viruses related to the family-level group Pimascovirales_01 (PM_01), providing the first host assignment for this poorly characterized viral lineage. Thus, scDNA-seq captured a broad landscape of potential host-virus associations, including lineages not detected by scRNA-seq, whereas viral transcriptomics identified interactions occurring during active infection of native cells. Together, these complementary approaches distinguished the broader landscape of viral associations from transcriptionally active infections, which could then be interrogated for viral activity and host responses. Notably, however, dinoflagellates were detected by scRNA-seq but were absent from scDNA-seq. This difference persisted among uninfected cells (fig. S2) and contrasted with their high abundance in the bulk metagenomes (Fig. 2A), suggesting that their absence from scDNA-seq may reflect differences in sample handling or sequencing methodology.

### High diversity and specialization of giant viruses from Gossenköllesee

To place the giant viruses discovered in Gossenköllesee in a taxonomic context, we reconstructed a phylogeny of GVMAGs together with genus-level representatives from the Giant Virus Database (GVDB; Fig. 3, data S6). Gossenköllesee GVMAGs were distributed across most major giant virus orders, highlighting the high diversity of giant viruses in this lake. Several metagenome-derived giant viruses were detected in single cells (Fig. 3), although not all could be confidently assigned to a host. Although multiple distinct viruses from the candidate order “Pandoravirales” were identified via scDNA-seq, none directly matched the specific pandoravirad GVMAGs assembled from bulk metagenomic data (Fig. 3). This suggests that the lake harbors distinct, coexisting pandoravirad lineages that are differentially captured by bulk and single-cell sampling techniques. This highlights the complementarity of the two approaches, with scDNA-seq capturing viral lineages not recovered by bulk metagenomic sequencing and assembly. A similar pattern has been observed in marine viral communities: Vila-Nistal et al. (*27*) found that <1% of species and genus-level viral clusters were recovered across all sequencing approaches tested, with single-virus genomics yielding a particularly distinct taxonomic profile. Only a small subset of the viral lineages was detected by scRNA-seq, consistent with active viral transcription occurring in only a fraction of cells. These phylogenetically resolved genomes provided the reference framework for characterizing viral transcription in actively infected cells.

**Fig. 3.**
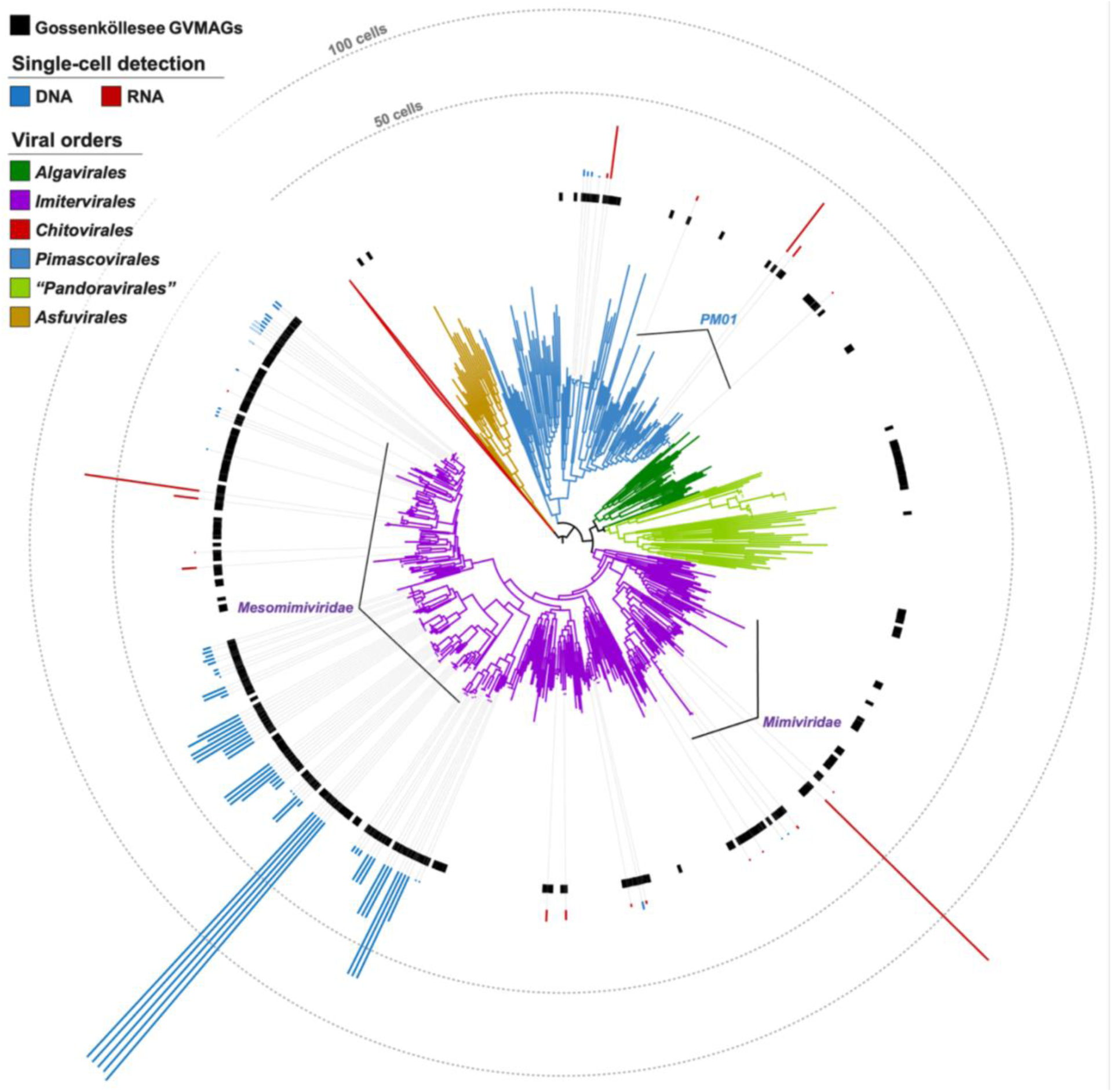
Phylogenetic affiliation of Gossenköllesee giant virus genomes recovered by bulk metagenomics. Branch colors indicate viral orders. The black ring denotes the Gossenköllesee GVMAGs. The lengths of the red and blue bars indicate the number of single cells in which each virus was detected by scDNA-seq and scRNA-seq, respectively. Dashed circles indicate 50 and 100 cells.

### An unusual viral TIR-nuclease gene is expressed during chrysophyte infection

To characterize viral genes expressed during active giant virus infection, we mapped the assembled single-cell contigs from infected cells back against the bulk GVMAG catalog. We identified three dominant host-virus systems undergoing active viral transcription (fig. S4, fig. S5A to C, data S7). Among these, 40 chrysophyte cells were infected by the mesomimivirid AIL3.4. From the genes expressed by AIL3.4, we identified a protein containing a Toll/Interleukin-1 receptor (TIR) domain (pfam13676) fused to a YncB nuclease domain (COG1525) (fig. S5A and D, data S8). TIR domains have conserved roles in immune and cell-death pathways in bacteria, plants, and animals (*9*, *28*). TIR-containing proteins have been reported from viruses that infect haptophytes of the genera *Haptolina* and *Prymnesium* (*26*), but not in combination with a nuclease domain. We detected proteins containing both the TIR domain and the YncB nuclease in only 8 bacterial genomes from the Burkholderiaceae family, but with the domains in opposite order and lacking sequence homology to the viral protein (data S8). However, we did identify homologous proteins with the same TIR-nuclease architecture in three additional Gossenköllesee giant viruses and two viruses from an Arctic freshwater lake in Canada (*29*; JGI project IDs: Ga0370460 and Ga0335110). Co-option of host defense mechanisms by viruses has previously been described in bacteriophages; for example, the anti-Pycsar protein Apyc1 counteracts bacterial cyclic nucleotide signaling defenses (*30*). This raises the possibility that the viral TIR-nuclease protein may represent an anti-defense strategy. In addition, its conservation among giant viruses from geographically distant cold freshwater lakes suggests that it represents an ecologically relevant adaptation.

### Dinoflagellates induce a conserved antiviral gene during natural giant-virus infection

Having resolved viral transcription within native host cells, we next asked whether we could detect host responses to infection in the same natural population. We focused on dinoflagellates infected by AIL3.53, an aliimimivirid related to *Cafeteria roenbergensis* virus. This host-virus pair represented the most prevalent active infection in our single-cell transcriptomic dataset. Family-level taxonomy could be assigned to 85% of Dinophyceae cells (fig. S6), and among these, 92% were classified as belonging to the family Thoracosphaeraceae, within the order Peridiniales (fig. S6). Thoracosphaeraceae comprise a diverse group of heterotrophic, photosynthetic, and parasitic dinoflagellates inhabiting both marine and freshwater environments (*31*, *32*). The Gossenköllesee Thoracosphaeraceae could not be confidently classified beyond the family level, potentially reflecting limited representation in reference databases or a hitherto uncharacterized lineage. Differential gene expression analysis comparing infected (≥10 viral UMIs) and uninfected dinoflagellate cells identified 72 significantly differentially expressed host genes (adjusted P < 0.05; Fig. 4, data S9). Among these genes, a subset exhibited a binary expression pattern, namely, transcripts were undetectable in uninfected cells but strongly induced during viral infection (Fig. 4A).

**Fig. 4.**
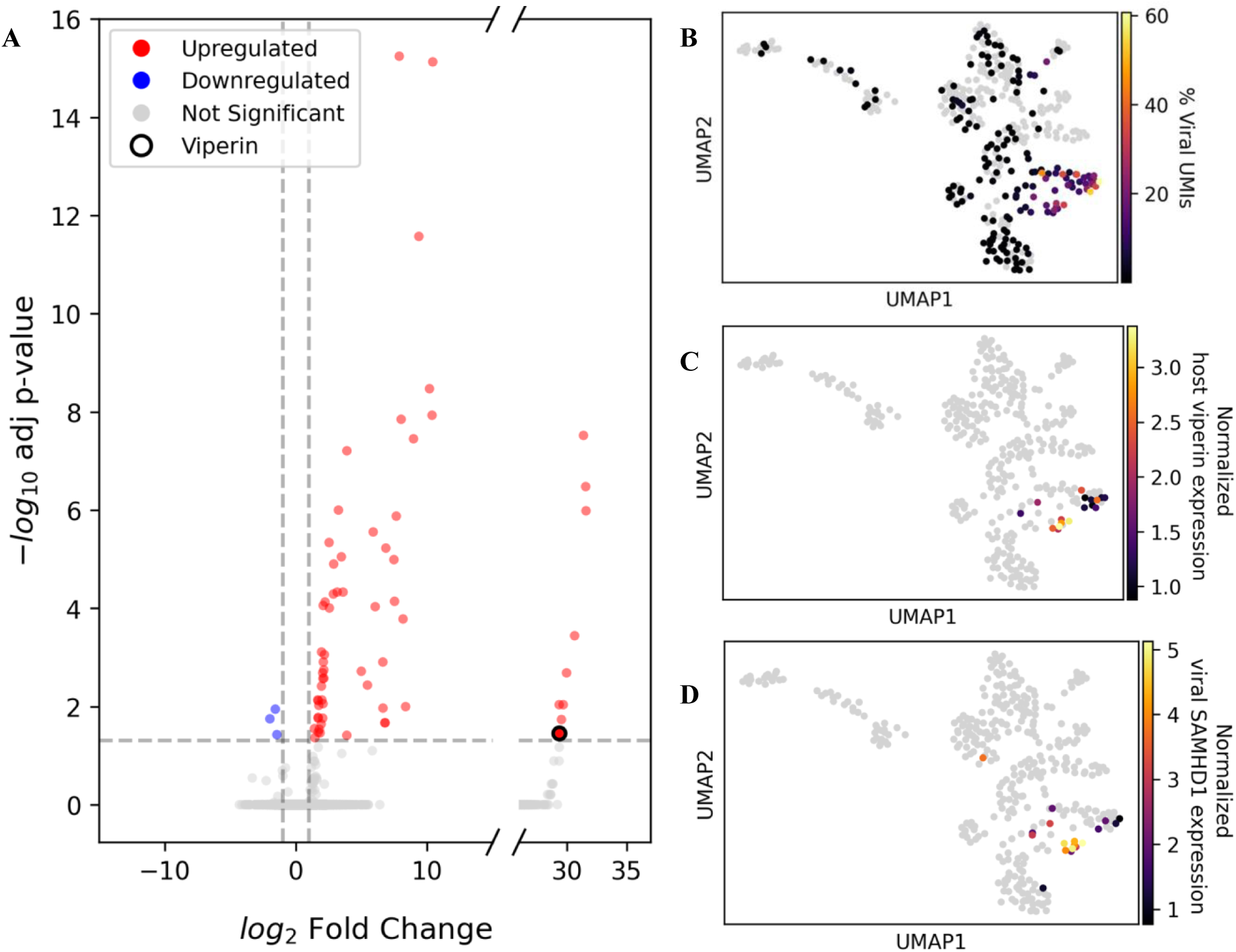
Expression of immunity-associated genes during giant-virus infection. (A) Volcano plot of host genes differentially expressed between infected (expressing ≥ 10 viral UMIs, log2 FC>1, adjusted *P* < 0.05) and uninfected cells. Statistical significance (-log10 adjusted P-value) is shown as a function of the log2FC. Dots in the right-hand panel (log2FC>25) represent infection-specific upregulated transcripts. These genes were undetected in uninfected cells, resulting in discrete, high-magnitude fold changes. (B-D) UMAP visualization of scRNA-seq data of dinoflagellates and viral transcription across individual cells (372 cells; 61 infected). Zero values are shown in gray. (B) Percentage of total UMIs assigned to viral transcripts in each cell. (C) Normalized expression of the dinoflagellate host viperin gene. (D) Normalized expression of the viral SAMHD1 gene.

Strikingly, among the infection-induced genes was a homolog of viperin, a member of an ancient family of antiviral genes conserved across bacteria and eukaryotes (*33*). Viperins are radical SAM enzymes that generate modified nucleotides capable of interfering with viral replication (*1*, *34*, *35*). The dinoflagellate protein retained the conserved catalytic motif required for viperin activity, and structural modeling revealed strong similarity to mouse viperin with a TM score of 0.91 (Fig. 5A). Within the natural population, viperin expression was restricted to infected cells and detected in 17 of 61 infected dinoflagellates (Fig. 4B, C). Thus, rather than merely encoding a conserved antiviral system, these protists selectively induced it during ongoing viral infection in the wild. This represents the first *in situ* detection of a conserved antiviral defense protein actively expressed by its single-celled eukaryote during viral infection.

**Fig. 5.**
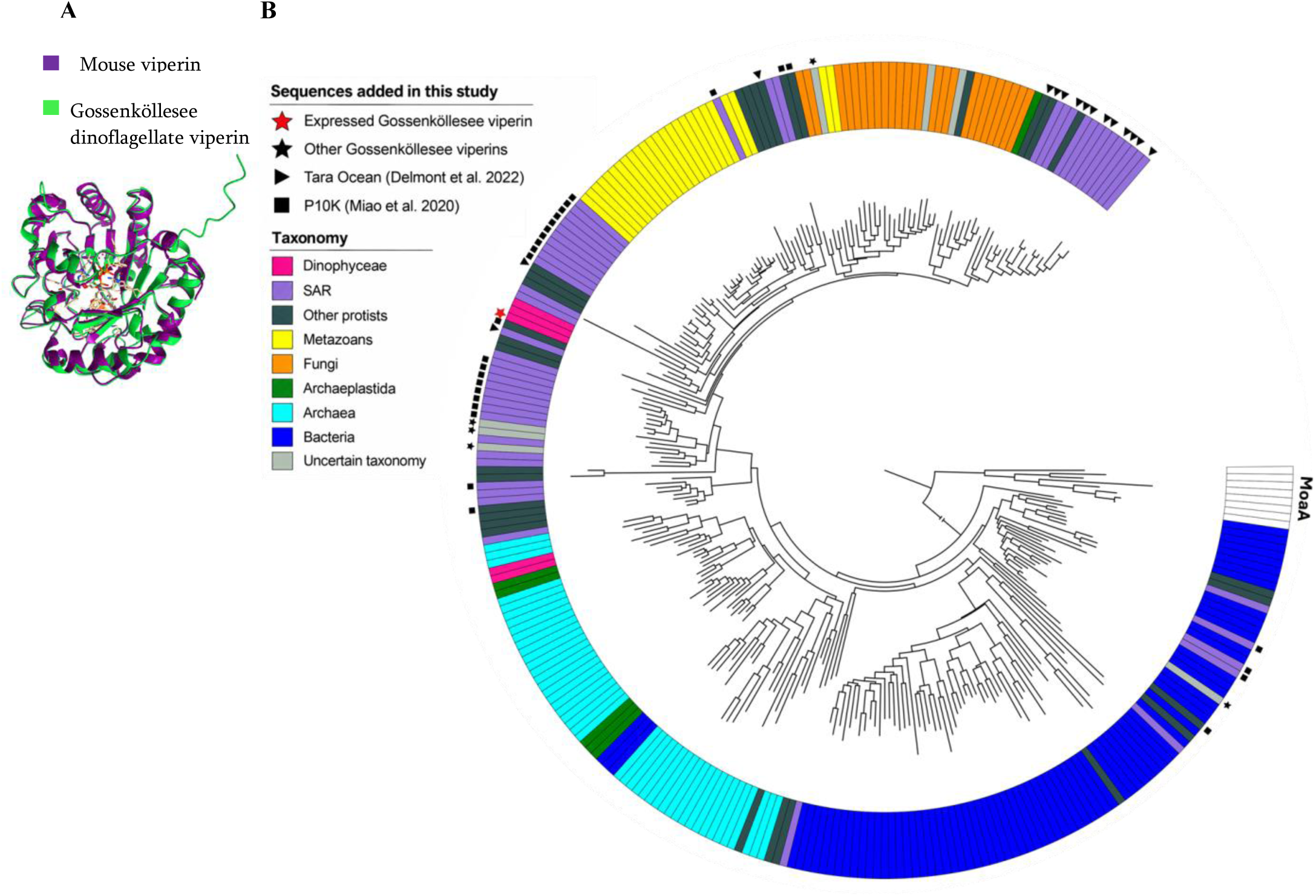
Gossenköllesee dinoflagellate viperin is part of a conserved antiviral protein family. (A) Predicted structure of Gossenköllesee dinoflagellate viperin (green) aligned with the crystal structure of mouse viperin (PDB: 6Q2Q; purple) bound to uridine triphosphate and S-adenosylhomocysteine. The N-termini of both structures are hidden. (B) Phylogenetic tree of viperins rooted using MoaA as the outgroup. Colors indicate taxonomic classification. Most sequences are derived from a previous study on viperin conservation (*33*), with additional sequences identified during this study or obtained from Tara Oceans (*38*) and from the P10K project (*39*). SAR: stramenopiles, alveolates, rhizarians.

### Viperin induction coincides with expression of viral immunity-associated genes

Viperin-expressing cells spanned a broad range of infection levels, with viral transcripts comprising 2 to 60% of total UMIs per cell (Fig. 4B, C). We searched among the viral transcripts for potential countermeasures against host immunity. Among the viral genes positively correlated with viperin expression (Spearman correlation ≥0.4, FDR <0.05) were homologs of a DEAD/DEAH-box helicase and SAMHD1, both associated with antiviral functions (Fig. 4C and D, fig. S7, data S9). SAMHD1 was shown to prevent reverse transcription of HIV-1 RNA through dNTP depletion (*36*) and DEAD/DEAH-box helicases are ATP-dependent RNA helicases that remodel RNA and participate in a variety of cellular functions, including cellular stress responses and antiviral immunity (*37*). Interestingly, the host itself also encodes for the SAMHD1 protein (fig. S8). The viral protein showed structural similarity to the host homolog, with a TM-score of 0.69 when normalized to the host protein length; normalization to the shorter viral protein yielded a TM-score of 0.87, indicating that most of the viral structure is contained within the host fold (fig. S8A). Host and viral SAMHD1 genes are co-expressed in some cells (fig. S8B), suggesting they may interact during infection. Together with our observation of a viral TIR domain-containing gene in a chrysophyte-infecting virus (fig. S5A, C), this suggests that viruses can utilize immunity-associated proteins as an anti-defense strategy. However, whether the viral homologs retain their canonical functions or have been repurposed for other roles in the viral life cycle remains unknown and requires experimental validation.

### Viperin is a conserved component of protist immunity

To place the dinoflagellate viperin in an evolutionary context, we expanded the viperin dataset from a previous study on viperin conservation across the tree of life (*33*), with homologs recovered from Gossenköllesee metagenomes, global ocean metagenomes (*38*), and aquatic protist cultures (*39*) and reconstructed the phylogeny of the expanded dataset (Fig. 5B; data S10). The expressed viperin in Gossenköllesee clustered with dinoflagellate and other SAR-associated homologs. It is also evident that fungal and metazoan viperins are nested within a predominantly protist viperin clade. Together with the widespread occurrence of viperin homologs in aquatic protists, this phylogenetic distribution suggests that viperin is an ancient and broadly conserved component of protist antiviral immunity. The widespread distribution of viperin among aquatic protists and its induction during natural giant-virus infection advance our understanding of how antiviral defense systems evolved during the transition from prokaryotes to multicellular eukaryotes. Importantly, our data provide an ecological dimension to this evolutionary conservation; rather than merely detecting the genomic potential for antiviral defense, single-cell RNA-seq captured the selective induction of a conserved antiviral system in its native host during ongoing viral infection in nature. Our results provide a direct link between viral infection and conserved antiviral immunity in a free-living protist population and extend the antiviral role of viperin from RNA (*40*, *41*) and DNA viruses (*42*) to giant viruses for which no defense system has been reported to date.

## Conclusions

The discovery of antiviral defense systems in bacteria and eukaryotes has transformed our understanding of the evolution of immunity. Here, by simultaneously resolving viral and host transcription within individual cells, we captured a conserved antiviral response during ongoing infection in a natural protist population. Our findings provide ecological context and molecular insights for some of the myriad host-virus interactions that shape protist communities in alpine lakes. In addition to recovering native host-virus pairs and deciphering viral gene expression in situ, we discovered an ancient antiviral defense system that is induced in protists during viral infection. This demonstrates the value of studying host-virus interactions directly in natural populations at single-cell resolution, where antiviral responses and viral infection strategies can be observed in their cellular and ecological context. Understanding the evolution of these systems and their widespread distribution among protists will be essential for reconstructing the evolutionary history of immunity across the tree of life.

## Materials and Methods

### Environmental sampling

Samples were collected from Gossenköllesee, an alpine lake in Austria (47°13′46″N, 11°00′50″E), during multiple sampling campaigns between November 2021 and August 2025 (table S1). An initial single-cell RNA-sequencing sampling campaign was conducted in August 2023. Five samples were collected, pre-filtered through a 55 μm mesh, and 2 L of sample water from each sample were collected on a 3 μm filter until the filter clogged. Samples were fixed by mixing 100 μL of concentrated sample with 900 μL of chilled methanol, then immediately placing them on dry ice. Samples were later stored at −80°C until library preparation. Because this initial campaign yielded relatively few cells, all five samples were pooled before library preparation.

The primary sampling campaign was conducted in August 2024, during which companion bulk metagenomic, single-cell DNA-seq, and single-cell RNA-seq datasets were generated. Four bulk metagenomic samples (APESAF, AIL3, TPESL, and TIL3J) were collected from composite (1-8 m) and deep (7-8 m) water samples, representing paired 0.22-40 μm and 3-40 μm size fractions (table S1). The sampling design encompassed the surface layer receiving full irradiance, the middle of the photic zone, and the lowest light intensity just above the lake bottom, where care was taken to avoid sediment. Water was pre-filtered twice through a 30 μm nylon mesh before 2 L were collected onto either 0.22 μm polyethersulfone or 3 μm polycarbonate filters. Filters were immediately frozen at −20°C until DNA extraction. For single-cell DNA sequencing, environmental water was pre-filtered to remove large particulate matter, gently concentrated by filtration, and stored at 4°C for 30 days before encapsulation. For single-cell RNA sequencing, two samples were pre-filtered multiple times through a 40 μm mesh. 2 L of sample water from each sample was collected on a 3 μm filter until the filter became clogged. The suspension was concentrated, and 100 μL of each sample pellet was fixed in chilled methanol, transported on dry ice, and stored at −80°C until library preparation.

To characterize seasonal variation in giant virus diversity, a monthly 5 μm metagenomic time series was collected between September 2024 and August 2025, excluding February due to thin ice and May due to adverse weather. Three water samples were collected around midday from a fixed platform at depths of 0, 4, and 8 m using a 5-L water-sample bailer (Uwitec). Samples were transported under light protection to the laboratory within 4 h of collection. For metagenomic analyses, 1 L of water from each depth was pre-filtered through a 100 μm plankton net and collected onto a 47 mm, 5 μm polyethersulfone filter (Delvstlab, WKLMQS-PES-50-5), which was stored at −80°C until DNA extraction. Finally, ten previously generated bulk metagenomic datasets (>0.22 μm size fraction) collected between November 2021 and November 2022 were included for giant-virus identification (*43*) (NCBI BioProject PRJNA1434480). Details of all samples, sequencing approaches, and downstream applications are provided in table S1.

### Bulk metagenomic sequencing

Four Illumina metagenomic datasets (APESAF, AIL3, TPESL, and TIL3J; table S1) were generated from the August 2024 sampling campaign. DNA was extracted using the DNeasy PowerWater Kit (Qiagen). Sequencing was performed by Novogene using Illumina paired-end sequencing (2 × 150 bp), targeting approximately 60 Gb per sample. Sequence data were deposited under NCBI BioProject PRJNA1483949.

### 5 μm time-series metagenomes

DNA was extracted from the 5 μm filters using the DNeasy PowerWater Kit (Qiagen) and sequenced using Illumina paired-end sequencing (2 × 150 bp), targeting approximately 20 Gb per sampled depth (60 Gb per time point). Libraries generated from the three depths at each time point were pooled before metagenomic assembly.

### 0.22 μm time-series metagenomes

Ten previously generated metagenomic datasets (>0.22 μm size fraction) collected between November 2021 and November 2022 (*43*) (NCBI BioProject PRJNA1434480) were included for giant virus identification (samples GKS0-GKS9; table S1).

### Assembly and binning of Metagenomic sequencing data

For each sample (GKS0-GKS9-5µm, APESAF, AIL3, TPESL, TIL3J; Supplementary table S1), raw reads were trimmed for Illumina adapters using Trimmomatic v0.39 (*44*). Trimmed reads from APESAF, AIL3, TPESL, and TIL3J were assembled using SPAdes (*45*) using parameters: --meta --no-error-correction -k 23,33,43,53,63,73. In the case of the 5 µm time-series (GKS0-GKS9-5µm), the three sets of reads from 0, 4, and 8 m depth at each time point were pooled and assembled using MEGAHIT v1.2.9 (*46*) (PMID: 25609793) with the flags --k-list 23,33,55,73 --min-count 3 --presets meta-large, creating nine metagenomic assemblies. From the 2021-2022 time series, (GKS0-9-0.22µm) contigs >5 kb from each assembly were binned separately using MetaBAT2 (*47*). Differential coverage tracking across all 10 time-series metagenomes was utilized to improve binning resolution, enforcing a minimum bin size of 100 kb. Generated bins were then de-replicated using dRep at 95% Average Nucleotide Identity (ANI) (*48*). Bins over 100 kb in length were retained using the following execution parameters: -l 100000 --ignoreGenomeQuality --S_algorithm ANImf --P_ani 0.9 --S_ani 0.95 --cov_thresh 0.1 --coverage_method large. Bulk metagenomic diversity was assessed using phyloFlash (*49*).

### Creating a reference gene catalog from the metagenome

To create a reference gene catalog for the microbial communities in Gossenköllesee, contigs longer than 5000 bp from each sample were used to predict genes using Genemark-ES v. 4.72 (*50*), and Metaeuk v. 7 (*51*). These programs also generated a set of translated proteins from the predicted genes. Genes and proteins predicted from both programs were combined into a single FASTA file. Genes were dereplicated with 0.95 identity and a minimum coverage of 200 bp using MMseqs2 (*52*). De-replicated genes were dereplicated again with an identity threshold of 0.99 using CD-HIT v. 4.8.1 to remove redundant contigs shorter than 200 bp (*53*). The final set of proteins was based on the twice-replicated gene set.

### Reference gene catalog annotation

Functional annotation of the gene catalog was performed on the final set of translated proteins using emapper v.2.1.13 (*54*) and Diamond v.2.1.16 (*55*), based on the eggNOG database v.5.0.2 (*56*). Conserved domains were identified on the final set of translated proteins using HMMER 3.4 (*57*). To identify viral genes in the gene catalog, predicted genes were compared against a custom protein database that combined RefSeq v. 207 (*58*) with all predicted proteins in the Giant Virus Database (GVDB) (*61*) (constructed as described above) using a translated alignment approach. Predicted genes were compared to the constructed database using LASTAL v. 959 (*59*) (parameters -m 100, -F 15, -u 2) with best matches retained.

### MAG generation and screening

To identify MAGs that encoded NCLDV (GVMAGs), we used Viral Recall v. 2.0, removing contigs that scored < 0, which are potentially non-viral (*60*), using the -c option. This resulted in 272 GVMAGs across all datasets combined. We then constructed a phylogenetic tree of these GVMAGs together with genus-level representatives chosen from the Giant Virus Database (GVDB) (*61*). For this, we used the ncldv_markersearch tool, which identifies 7 viral marker genes in the GVMAGs. NCLDV marker genes from all viral bins were translated and combined into a single protein FASTA file. They were clustered using CD-HIT v. 4.8.1 at 99% amino acid identity to remove redundancy (*53*). A LASTAL database was built from the gene catalog using LASTAL v. 959 (*59*).

### Single-cell DNA sequencing of eukaryotic microorganisms

Individual eukaryotic cells were isolated using the Semi-Permeable Capsule (SPC)-based high-throughput single-cell genomics workflow (Atrandi Biosciences®). Cell suspensions were encapsulated using an aqueous two-phase microfluidic system to generate individual semi-permeable capsules (SPCs). Half of the produced SPCs with isolated samples were treated with a PMA dye to evaluate the proportion of free-floating DNA encapsulated. Then, after obtaining an approximate sample concentration (after determining lambda values), two additional encapsulations were performed with differently diluted samples to obtain a suitable lambda value for sample barcoding and sequencing library preparation. Whole-genome amplification (WGA) of individual cells was performed using multiple displacement amplification (MDA) with WGA polymerase, following the manufacturer’s recommendations. Amplified DNA products were then quantified using Qubit 1X dsDNA HS Assay Kit (Invitrogen). The libraries were then sequenced on the MiSeq platform using the MiSeq Nano Kit v2 (300-cycles). Sequencing adapters and quality filtering were performed using Trimmomatic (*44*), with a minimum quality score of 20, and assembled de novo using SPAdes (*45*) in metagenomic mode.

### Eukaryotic Diversity Assessment in Single-Cell Genomics

Taxonomic affiliation was performed using a consensus approach among three methods: SAGs were taxonomically classified using 18S rRNA gene identification with Barrnap (*62*) and subsequently aligned against the SILVA database v138.2 (*63*) using BLAST (*64*). Secondly, similarity searches for each contig within the SAGs were performed against the NCBI RefSeq database using MMseqs (*52*). Thirdly, the overall taxonomic affiliation of SAGs was calculated with the software GVclass (*65*). Finally, consensus was calculated to the last common ancestor of the three methods or classified as unknown and/or chimeric when a lack of classification or conflicting classifications were observed.

### Identification of giant virus sequences in single-cell genomics

To identify giant viruses, assembled metagenomic contigs from all SAGs were screened for hallmark Nucleocytoviricota genes, for this, protein prediction was performed using Prodigal (*66*), with the flags -n and -p meta; subsequently, hidden-Markov models (HMM) for marker genes including major capsid protein (MCP), family B DNA polymerase (PolB), packaging ATPase, and viral transcription factor genes was performed using NCLDV_search tool (*61*). To identify further viral contigs lacking marker genes, candidate SAGs containing NCLDV sequences were examined using ViralRecall (*60*). Quality assessment and contamination detection of candidate viral sequences were performed with Gvclass (*65*). Final taxonomic assignment of giant virus metagenomic assembled genomes (gvMAGs) was performed using TIGTOG (*67*). Pairwise genome-wide average-nucleotide-identity (ANI) comparisons between gvMAGs were carried out using pyANI-plus (*68*) with the MUMmer algorithm (*69*) and plotted with the ggplots R package (*70*) (fig. S3).

### Library preparation and RNA-seq using 10X Genomics

For 10X Genomics analysis, samples were thawed and gently mixed, and 1.7 ml of each sample was transferred into Lobind DNA Eppendorf tubes and centrifuged at 3000 g for 3 min at 4 °C. The methanol mix was discarded and replaced by 400 µl of 1X PBS and 0.04% BSA. Cell suspension was filtered through a 40 μm cell strainer (Falcon), and cell concentration was measured using a CytoFLEX S Flow Cytometer (Beckman Coulter) based on forward and side scatter. Cell concentration ranged from ∼3000 cells µl^-1^ to 8000 cells µl^-1^. Because of the low cell abundance recovered from the 2023 sampling campaign, the five samples collected that year were pooled before library preparation to achieve the target recovery of 10,000 cells. All concentrations were brought to 1000 cells µl^-1^ to target 10,000-20,000 cell recovery, according to the 10X Genomics Cell Suspension Volume Calculator Table provided in the user guide. The cellular suspension was loaded onto Next GEM Chip G, targeting 10,000 cells (v3.1 Chemistry; 2023 sample) or 20,000 cells (v4 Chemistry; 2024 samples), and then run on a Chromium Controller instrument to generate GEM emulsion (10X Genomics). A single-cell 3’ RNA-seq library was generated according to the manufacturer’s protocol (10X Genomics Chromium Single Cell 3’ Reagent Kit User Guide v3.1 Chemistry for the 2023 sample; v4 for the 2024 samples), with 11-12 cycles for cDNA amplification and 16 cycles for library amplification. Library concentrations and quality were measured using the Qubit dsDNA High Sensitivity Assay kit (Life Technologies). Libraries were pooled based on the target cell number, aiming for at least 20,000 reads per cell. Pooled libraries were sequenced using the Illumina NovaSeq X (100 cycles).

### Mapping scRNA-seq reads against the gene catalog

A pseudo-GTF file for the combined database was created using Bioawk v. 11 (github.com/lh3/bioawk). A reference was created from the database using the Cell Ranger mkref command. The Cell Ranger Software Suite (v. 10.0.0) was used to perform barcode processing (demultiplexing) and single-cell unique molecular identifier (UMI) counting on the raw reads using the count script (default parameters), with the deduplicated gene catalog database as a reference. All barcodes within the Cell Ranger-filtered matrix were used for downstream analysis.

### Identifying the taxonomy of individual cells by sequence homology to ribosomal RNA

Raw reads from each cell were pulled by the cell’s unique barcode identifier using seqtk v. 1.2 (https://github.com/lh3/seqtk). Reads were then trimmed (command: trim_galore --phred33 -j 8 --length 36 -q 5 --stringency 1 --fastqc -e 0.1), and poly-A was removed (command: trim_galore --polyA -j 1 --length 36), using TrimGalore (v. 0.6.5), a Cutadapt wrapper (*71*). Trimmed reads from each cell were assembled using RNASPAdes 3.15 (*45*) with kmer 21, 33, and 55. Raw reads were pulled, trimmed, and assembled using an in-house script. To identify the cell taxonomy, assembled contigs from each cell were matched against 18S rRNA sequences from the Protist Ribosomal Reference (PR2) (*72*) and metaPR2 (*73*). To remove redundancy, the sequences in each database were clustered using CD-HIT v. 4.8.1 at 99% identity (*53*). 18S rRNA contigs were filtered using Barrnap (*62*) with the --euk option and then aligned to the PR2 and metaPR2 databases using BLASTN (*74*), at 98% identity, E-value ≤ 10^-10^ and alignment length of at least 300 bp. Contigs were ranked by their bit scores, and only the best hits were retained for each contig. Contigs that matched more than one of these taxonomic groups were considered non-specific and were therefore ignored. Chimeric contigs were defined as contigs containing regions that match different taxonomic groups. Cells with chimeric contigs were also excluded to avoid doublets. To avoid detection of doublets and predators, cells that transcribe 18S rRNA transcripts homologous to more than one taxonomic group were conservatively omitted. Of the 7,127 cells detected, 3,419 could be identified using assembled 18S rRNA transcripts, and 3,708 (52%) were omitted because we could not assemble specific 18S rRNA contigs from them or because their identities were ambiguous.

### Identifying the infecting virus using a homology search against a custom protein database

To identify transcripts derived from giant viruses, assembled contigs from detected cells were compared to the reference NCLDV gene catalog protein database using a translated alignment approach using LASTAL v. 959 (*59*) (parameters -m 100, -F 15, -u 2). To verify that suspected viral contigs are viral, they were compared to a database combining RefSeq v. 207 (*58*) with all predicted proteins in the Giant Virus Database, constructed as described above. Contigs that matched non-viral organisms were removed, and the final contig set was analyzed at different taxonomic levels, consistent with the Giant Virus Database (for giant viruses) or NCBI taxonomy (for everything else) (*75*). Of the 3,419 taxonomically identified cells, 115 were classified as infected if they contained at least two assembled viral transcripts assigned to the same viral family, with ≥90% amino acid identity and an alignment length of at least 30 amino acids (Fig. 2). Only 91 cells with at least 90% of viral transcripts matching the same family are shown on the graph. Cells identified as rotifers were removed because they likely represented contamination resulting from fragmented microscopic animals passing through the filters. A Sankey plot was constructed using Holoviews v. 1.15.4.

### Phylogenetic analysis of GVMAGs

Viral marker genes in the GVMAGs were aligned using Clustal Omega, as in previously reported methods (*22*). We trimmed the alignment with trimAl (-gt 0.1) (*76*), generated the tree with IQ-TREE v. 2.2.2.7 (model LG+F+R10, 1000 ultrafast bootstraps) (*77*), and visualized it using the interactive Tree of Life (iTOL) (*78*).

### Comparison of GVMAGs with single-cell datasets

To examine how many SAGs corresponded to GVMAGs, we used the ANIcalculator tool, with default parameters (*79*). We used SAGs as the query, GVMAGs as the reference, and considered only matches with >95% nucleotide identity. To identify cells in the sc-RNAseq dataset with matches to GVMAGs, we mapped all assembled scRNA-seq transcripts against a custom database that included all GVMAGs, SAGs, and a large number of reference GVMAGs compiled elsewhere (*29*), using lastal v. 959 with default parameters (*59*). We only considered matches with >95% identity and alignment lengths >20, and cell-virus pairs with >4 matches were retained to avoid false positives. We masked the custom nucleotide database and query with TANTAN v. 22 (*80*) (default parameters) before searching to limit matches based on repetitive sequences.

### Viral gene expression visualization

Single-cell RNA sequencing reads were aligned to each viral genome individually using STARsolo v2.7.11 (*81*). Genome indices were generated from the viral genome assemblies together with GTF annotation. Cell barcodes and unique molecular identifiers were extracted from Read 1 using the corresponding cell lists for each taxon infected by a specific virus. Read 2 was aligned to the viral reference using local alignment with adapter clipping enabled (--clipAdapterType CellRanger4). To reduce spurious alignments, only alignments satisfying a minimum aligned fraction of 90% of the read length (--outFilterMatchNminOverLread 0.9), a minimum alignment score of 90% of the read length (--outFilterScoreMinOverLread 0.9), and a maximum mismatch rate of 5% (--outFilterMismatchNoverLmax 0.05) were retained. PCR duplicates were removed using the Cell Ranger one-mismatch UMI collapsing algorithm (--soloUMIdedup 1MM_CR). Gene-level UMI count matrices were generated with the STARsolo Gene mode (--soloFeatures Gene) and imported into AnnData for downstream analyses. To visualize transcriptional activity across viral genomes, coding sequence annotations were combined with the corresponding genome assemblies to generate pseudo-genome coordinates preserving the complete genomic organization, including intergenic regions. EggNOG functional annotations were predicted for each virus (*56*). For fragmented assemblies, contigs were concatenated with fixed spacer regions while maintaining their original lengths. For each predicted coding sequence, the mean number of UMIs per expressing cell was calculated, and the number of cells expressing the gene was determined. Circular genome maps were generated in Python using Matplotlib. Bar position corresponded to the genomic coordinates of each coding sequence, scaled bar height represented the mean UMI count per expressing cell, and bar color represented the number of cells expressing the gene. Genes of particular interest were annotated with leader lines and labels positioned outside the genome track.

### Identification of viral TIRs

The viral TIR was identified using EggNOG (*56*). The nuclease domain was identified by the NCBI conserved domain database (CDD; e-value < 1e-10) (*82*). Additionally, bacterial genomes with a similar (but opposite) gene structure were also identified using the NCBI conserved domain database (CDD; e-value < 1e-10) (*82*), but did not share sequence similarity with the viral TIR-nuclease. To verify that the TIR-nuclease fusion was not an assembly artifact, scRNA-seq reads were mapped to the predicted TIR-nuclease locus using BWA v0.7 (*83*) and visualized using IGV (*84*). Other TIR-nuclease-containing proteins were identified from the Gossenköllesee gene set reference and a published set of metagenomic viral genomes (*29*) using BLAST homology (P-value < 0.05) and were verified using the NCBI conserved domain database (e-value < 0.001).

### Preprocessing of transcript abundance and dimensionality reduction

Preprocessing was performed using the Scanpy toolkit v.1.9.3 (*85*). The raw heterogeneous dataset of 7,127 cells expressing 108,653 genes was filtered to include only 70,282 genes expressed in at least 2 cells, using scanpy’s sc.pp.filter_genes with min_counts=2. The heterogeneous dataset was subset to include only Dinoflagellata (422 cells). The subset was filtered again to include only genes expressed in at least 2 cells. Cells with the lowest 1% number of UMIs, as compared with the distribution of transcripts per cell in the entire dataset, were removed for downstream analyses. To prevent doublet or multiplet cells, which can be biological (cell digestion) or technical (fused cells), cells with the highest 1% of UMIs were also removed. Mitochondrial and ribosomal genes were identified based on EggNOG functional annotations. The preprocessed dataset includes 372 cells and 7,248 genes. For each cell, the percentage of total counts attributed to mitochondrial genes was calculated. Cells with mitochondrial content exceeding the median plus five median absolute deviations (MADs), capped at 20%, were removed, as high mitochondrial expression is indicative of damaged or dying cells. After mitochondrial quality control, mitochondrial and ribosomal genes were removed from the dataset to prevent these gene classes from driving the embedding structure. To account for differences in sequencing depth across cells, raw counts were normalized per cell to a fixed library size of 10,000 before log transformation. Normalized counts were then log-transformed using a natural log (log1p) transformation. Log transformation metadata were explicitly stored to ensure compatibility with downstream analyses. Expression values were scaled to unit variance and zero mean, with extreme values clipped to reduce the influence of outliers.

Principal component analysis (PCA) was then performed using the ARPACK solver, retaining a fixed number of principal components. The k-nearest neighbor graph was constructed from the principal component representation of the data using Scanpy’s sc.pp.neighbors function. The graph was computed using the PCA embedding (use_rep=“X_pca”) with n = n_neighbors nearest neighbors for each cell. Two nonlinear dimensionality reduction methods were then applied for visualization. Uniform Manifold Approximation and Projection (UMAP) was computed using sc.tl.umap with a minimum distance of 0.8 and a spread parameter of 2 to control the balance between local and global structure in the embedding. T-distributed Stochastic Neighbor Embedding (t-SNE) was computed using sc.tl.tsne with perplexity = 60 and learning rate = 500. A fixed random seed (random_state = 42) was used for both algorithms to ensure reproducibility of the embeddings.

### Dinoflagellate differential expression analysis

To identify viral genes in the dinoflagellate expressed gene catalog, we combined EggNOG annotations, LASTAL-based viral annotations (see *Reference gene catalog annotation*), and nucleotide homology to the dinoflagellate-infecting virus AIL3.53. Homologs to AIL3.53 were identified using BLASTN with a minimum alignment length of 100 bp and nucleotide identity ≥90%. Differential gene expression analysis was performed using the Scanpy toolkit between infected and uninfected cells, with cells expressing at least 10 viral UMIs considered infected (*13*, *14*). Gene expression in infected cells was compared with that in uninfected cells using the Wilcoxon rank-sum test, using sc.tl.rank_genes_groups. The analysis was conducted on log-normalized expression values. For each gene, the Wilcoxon test was used to assess whether expression levels in infected cells differed significantly from those in the uninfected reference group. Resulting test statistics, P-values, and log fold changes were extracted into a tabular format using sc.get.rank_genes_groups_df. Genes were considered significantly differentially expressed if they passed a multiple-testing-adjusted *P* < 0.05.

### Structure prediction for viperin

The predicted sequence of the putative viperin was extended upstream by 24 amino acids to the first methionine residue. The structure of the putative viperin was predicted using AlphaFold v.2 via ColabFold (*86*). The viperin structural model was searched in the PDB database using Foldseek (*87*), which identified mouse viperin (PDB: 6Q2Q) as a close structural homolog (TM-score = 0.90). The structure of the putative viperin was visualized using ChimeraX v.1.6.1 (*88*).

### Identifying viral genes correlated with viperin

Host-virus co-expression correlations were computed from the Dinoflagellata single-cell RNA-seq dataset (372 cells; infection status assigned using a viral UMI threshold of ≥10 UMIs per cell, yielding 61 infected cells). All correlation tests were restricted to infected cells. Viral genes were defined as described above. Before testing, pairs were filtered to require a minimum detection of 5% of infected cells for both the host and viral gene. For each host-viral gene pair that passed these filters, Spearman correlation coefficients were calculated from log-normalized expression values across all infected cells. Two-sided P-values were adjusted for multiple testing using the Benjamini-Hochberg false discovery rate (FDR) procedure across all tested pairs. Significant associations were defined as FDR ≤ 0.05 and Spearman ρ ≥ 0.4. The number of co-expressing cells (double-positive count) was recorded for each pair and used in downstream visualization. Viral functional annotations were obtained from eggNOG-mapper and merged onto significant pairs. For viperin (140101_m_AIL3), significantly correlated viral genes were summarized in a lollipop plot showing Spearman ρ, FDR-adjusted significance (-log₁₀ FDR), and co-expression cell counts.

### Structure prediction for SAMHD1

SAMHD1 genes (for both virus and host) were identified using the eggNOG annotation (see above). The structures of the putative SAMHD1 proteins were predicted using AlphaFold v.2 via ColabFold (*86*). The initially predicted host SAMHD1 protein contained a long, unstructured C-terminal region with no detectable homologs in NCBI. We therefore used an alternatively spliced protein predicted from the same ORF, but lacking the terminal exons, for subsequent structural analyses. A structural model for both host and virus was searched in the PDB database using Foldseek (*87*), which identified human SAMHD1 (PDB: 4TO5) as a close structural homolog (TM-score with host = 0.84; TM-score with virus = 0.78). Both structures were aligned using TM-align (*89*), yielding TM-score = 0.69 (if normalized to the length of the host protein) and TM-score = 0.87 (if normalized by the length of the viral protein). Structural alignments of the putative SAMHD1 proteins with the human homolog were visualized using ChimeraX 1.6.1 (*88*).

### Constructing the viperin phylogenetic tree

The viperin sequence was curated manually from the original assembled contig in the metagenomic data and renamed 140101_m_AIL3_edited. A phylogenetic tree was constructed from protein sequences reported in a published study on Viperin conservation (*33*). To expand the repertoire of protist viperins, we searched for homologs in published metagenomic data (*38*) and in cultured protist genomes from the 10,000 Protist Genomes project (P10K) (*39*). We chose homologs that contained the conserved CXXXCXXC viperin motif and were not derived from animals. We also searched the Gossenköllesee metagenomic database for other viperin proteins using the Pfam conserved domain search and identified additional predicted viperins that contained the conserved CXXXCXXC motif. We constructed a phylogenetic tree with the additional protist viperins. The tree was constructed according to the method described in Shomar et al. (*33*). Viperin sequences were aligned using the MAFFT option L-INS-I (*90*) and trimmed using trimAl with option -gt 0.3 (*76*). Sequences with fewer than 50 aligned columns were removed using trimAl using the option -resoverlap 0.5 -seqoverlap 50. Phylogenetic analysis was performed using IQ-TREE v.2.1.3 (*77*) with an LG+C60+F+G4 mixture model, with ultrafast bootstrap values calculated using UFBoot2+NNI from 2,000 iterations. MoaA sequences were used as an outgroup. The phylogenetic tree was visualized using iTOL (*78*).

### Usage of AI

ChatGPT 5.6 (OpenAI) and Grammarly were used for language editing and grammar checking. ChatGPT 5.6 (OpenAI) and Grok 4.6 (xAI) were used to assist with code development. The authors reviewed and validated all AI-assisted content and take responsibility for the final manuscript and analyses.

## Acknowledgments

We thank Dr. Shifra Ben-Dor and Dr. Pedro Escudeiro for advice on the sequence and structure of the putative viperin.

## Funding

The Simons Foundation International grant No. 735079 (A.V.)

European Research Council (ERC) grant agreement No. 101125221 (M.G.F.)

The Austrian Science Fund grant No. I5824 (R.S.)

The Austrian Science Fund grant No. P-34620 (C.B.).

National Institutes of Health R35 grant No. 1R35GM147290 (F.O.A.).

Gossenköllesee belongs to the national and international long-term ecological research network (LTER-Austria, LTER Europe and ILTER) and GLEON, the Global Lake Ecological Observatory Network.

## Author contributions

Conceptualization: AF, MGF, AV

Methodology: AF, TAS, CB, FOA

Formal analysis: AF, TAS, CB, FOA

Investigation: AF, TAS, CB, TSS, DS, MES, NE, KK, RS, MGF

Visualization: AF, TAS, CB, FOA

Funding acquisition: MGF, RS, CB, FOA, AV

Supervision: AV, FOA, MGF

Writing - original draft: AF

Writing - review & editing: AF, TAS, CB, TSS, DS, RS, FOA, MGF, AV

## Data, code, and materials availability

The bioinformatic pipeline used to pair hosts and viruses in scRNA-seq data was consolidated into a single, reproducible pipeline, “Virus-Host-Pairing” (“ViHoP”). A GitHub repository is available at github.com/amir-fromm/ViHoP. The scripts used for data subsetting and preprocessing, and for generating the figures, are available in the repository at github.com/amir-fromm/ViHoP/downstream_scripts. A detailed README file is available at the repository.

Bioproject accession numbers for raw sequencing data are listed under table S1. GVMAGs assembled from Gossenköllesee were deposited to Zenodo:

DOI: 10.5281/zenodo.22834835.

The gene catalog assembled from Gossenköllesee was deposited on Zenodo: DOI:10.5281/zenodo.23059995.

No new biological materials were generated in this study.

