## Supplementary Materials for "Single-cell-resolved natural giant virus infections reveal conserved protist immunity"

### Supplementary Figures

a b

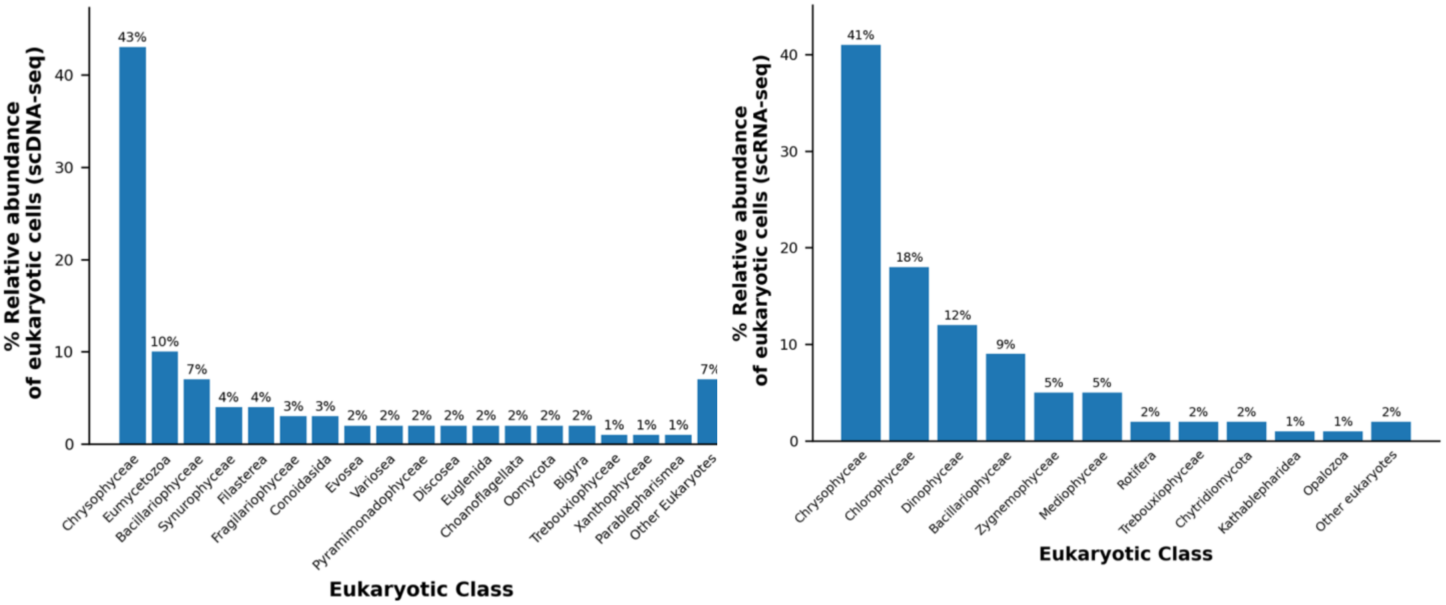


**Fig. S1. Eukaryotic community composition captured by single-cell sequencing.** (A) Relative abundance of major eukaryotic taxonomic groups recovered from ~4300 quality-filtered single-cell amplified genomes (SAGs) obtained from lake water samples. Relative abundances were calculated as the proportion of SAGs assigned to each taxonomic group within the eukaryote-classified SAGs subset, with the number on top of each bar representing the respective percentage. Groups with relative abundance <1% are displayed collectively as "Other Eukaryotes". (B) Relative abundance of eukaryotic classes from 3,419 cells sequenced by scRNA-seq, identified by 18S rRNA. Relative abundances were calculated as the proportion of cells assigned to each taxonomic group within the eukaryote-classified subset, with the number on top of each bar representing the respective percentage. Groups with relative abundance <1% are displayed collectively as "Other Eukaryotes".

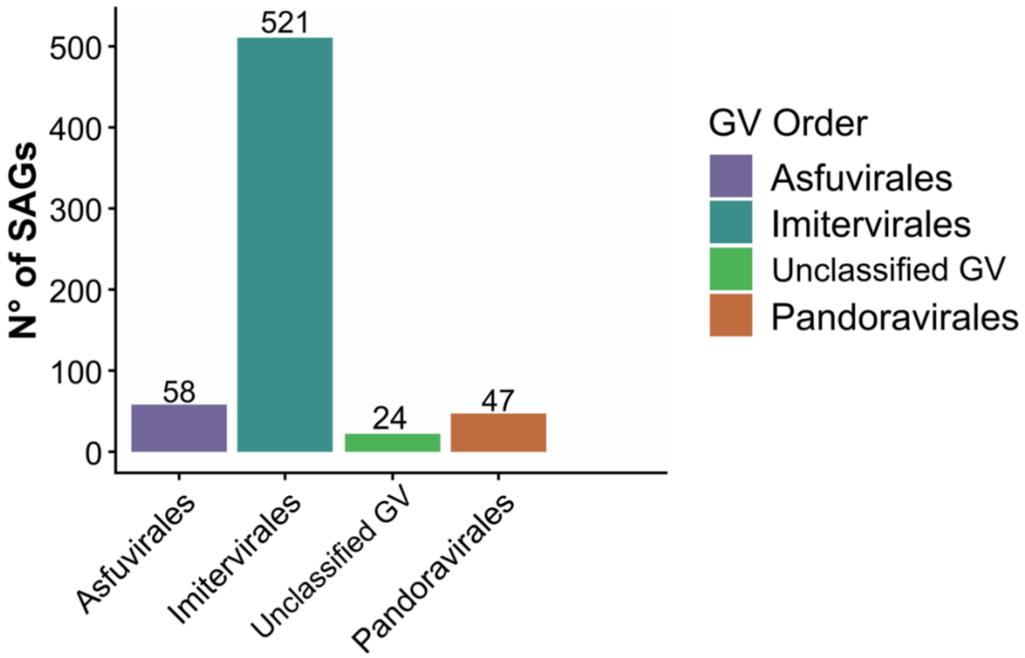


**Fig. S2. Composition of giant viruses in Gossenköllesee SAGs.** Relative abundance of Nucleocytoviricota lineages inferred from single-cell amplified genomes (SAGs). Viral genomes (vSAGs) were recovered using NCLDV marker genes and assigned to taxonomic orders using a machine learning-based approach implemented in TIGTOG. Imitervirales represented the dominant viral lineage, followed by Asfurvirales and Pandoravirales. Viruses with uncertain taxonomic classification are shown in the "Unclassified" category.

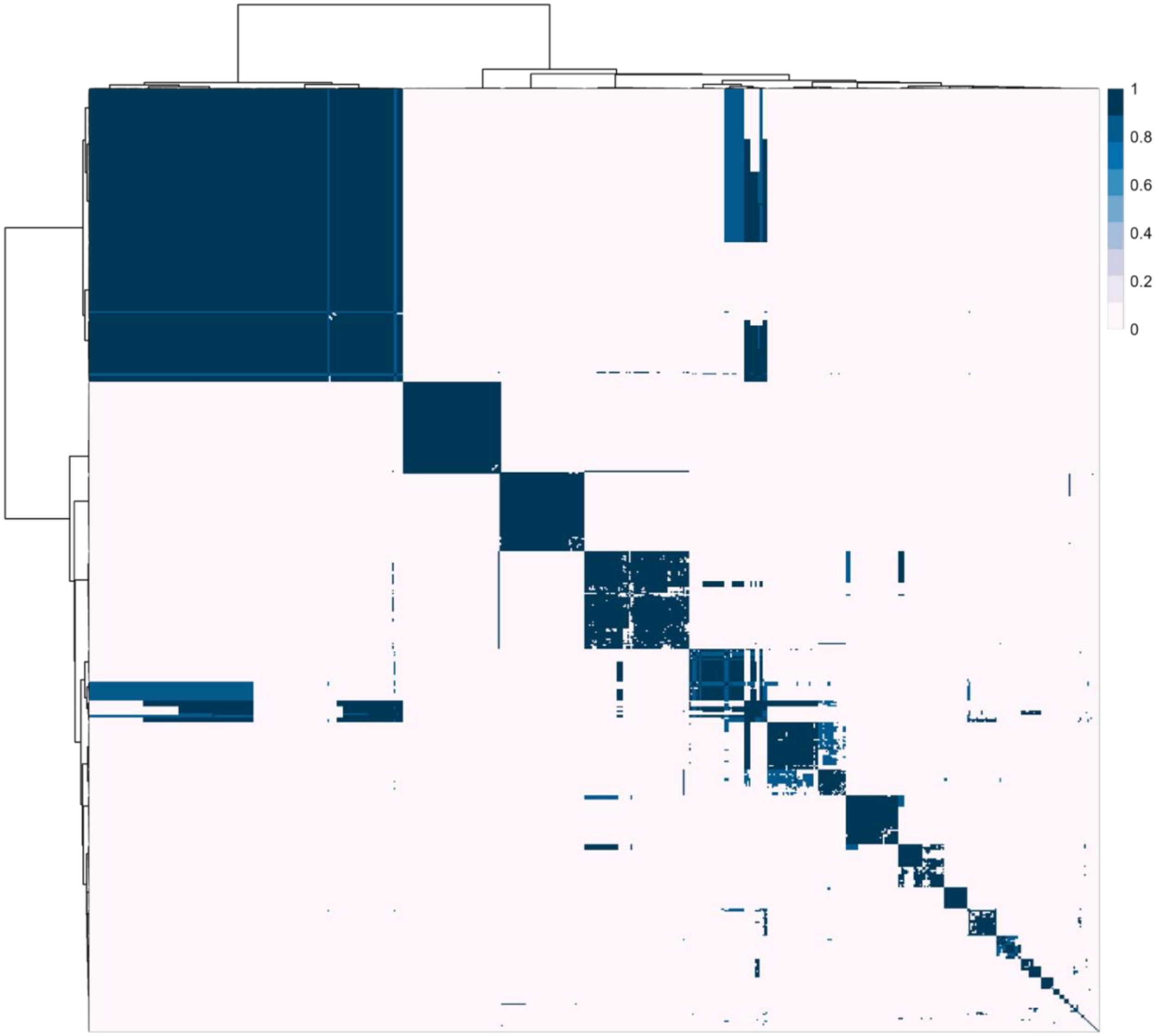


**Fig. S3. Pairwise average nucleotide identity (ANI) among Nucleocytoviricota genomes recovered from Lake Gossenköllesee SAGs.** Heatmap showing pairwise ANI values calculated between the 650 recovered viral metagenome-assembled genomes (vMAGs). Rows and columns represent individual vMAGs, ordered according to hierarchical clustering based on ANI similarity. Color intensity indicates nucleotide identity, with darker colors corresponding to higher ANI values.

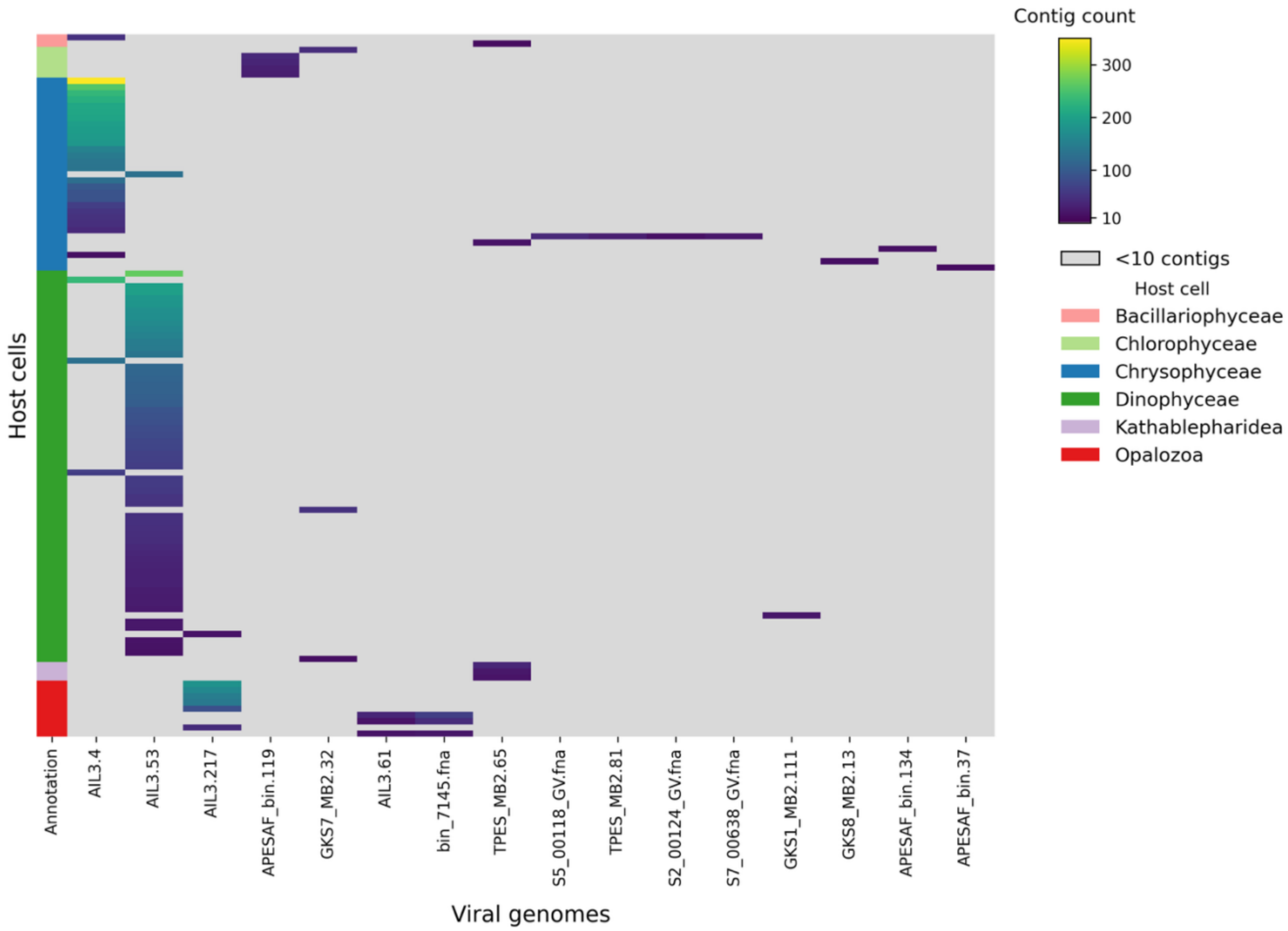


**Fig. S4. Nucleotide mapping of single-cell RNA-seq assembled contigs to full viral genomes.** Nucleotide sequences were mapped to the full genome of GVMAGs assembled from bulk metagenomics. Matches with fewer than 10 hits are greyed out.

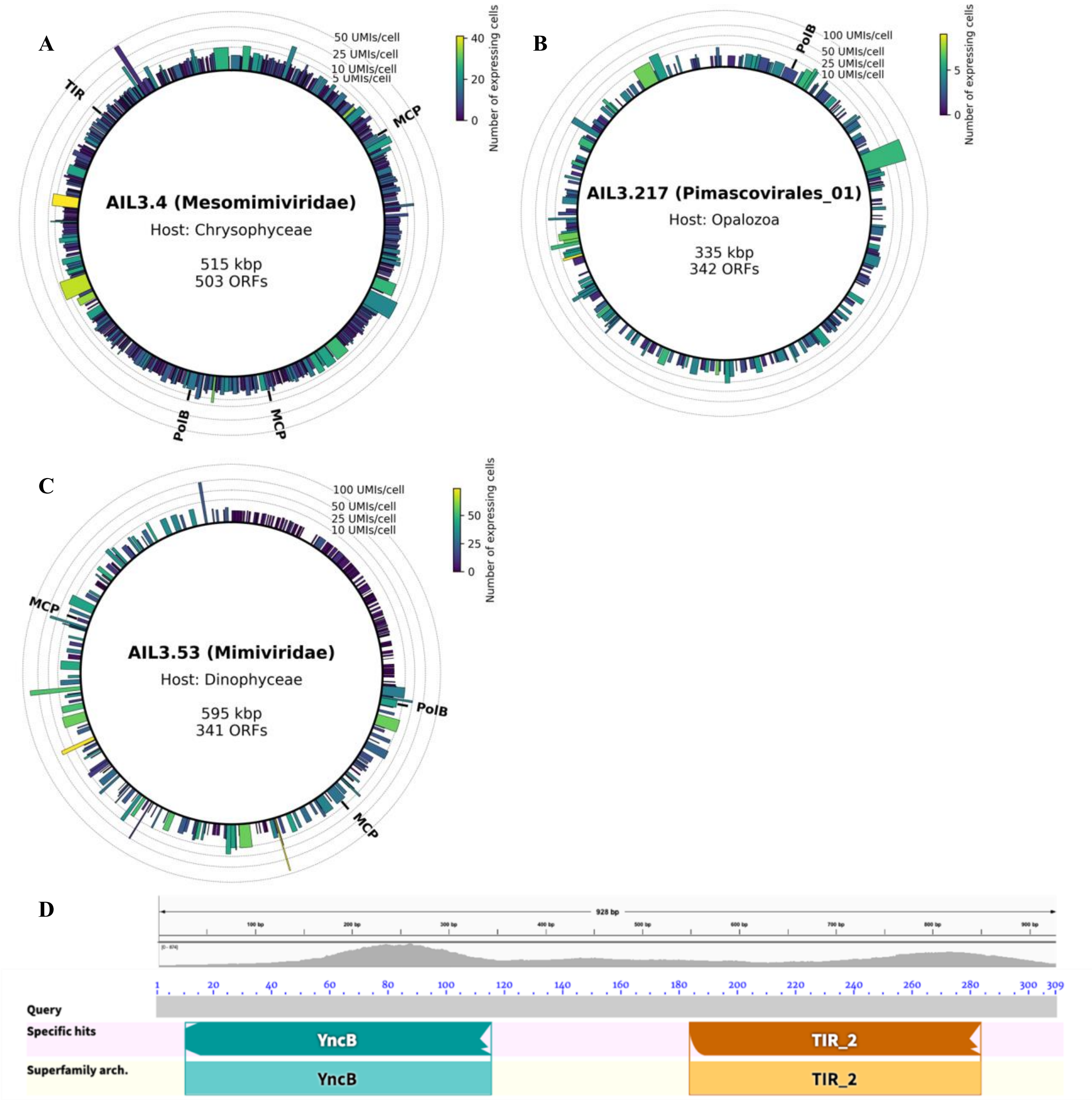


**Fig. S5. Genomics of giant viruses identified in Gossenköllesee.** (A-C) Genomic organization and transcriptional profiles of metagenome-assembled giant viruses detected in scRNA-seq data. Bar heights represent the mean number of Unique Molecular Identifiers (UMIs) per cell expressing the gene. Bar color is determined by the number of cells expressing the gene. Highlighted gene abbreviations: PolB: DNA Polymerase beta; TIR: Toll Interleukin Receptor-Nuclease; MCP: Major Capsid Protein. MCP was not identified in the AIL3.217 genome. (D) Gene organization of the putative viral TIR-like protein (bottom) and scRNA-seq read coverage.

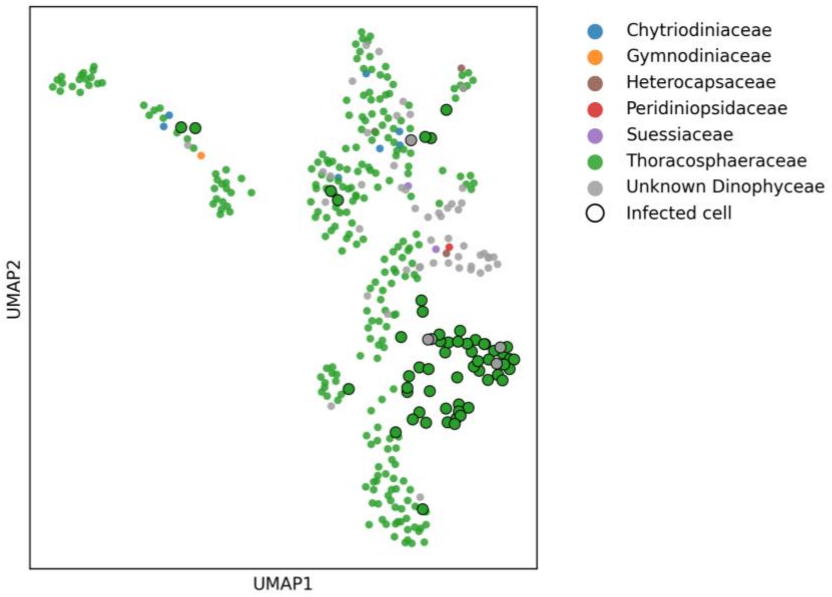


**Fig. S6. Taxonomic composition and infection status of dinoflagellates identified by scRNA-seq.** UMAP projection of dinoflagellate cells identified by scRNA-seq. Cells are colored according to their predicted family based on 18S rRNA homology (*n* = 372). Cells expressing at least 10 viral UMIs were classified as infected and are shown as enlarged points with black outlines (*n* = 61).

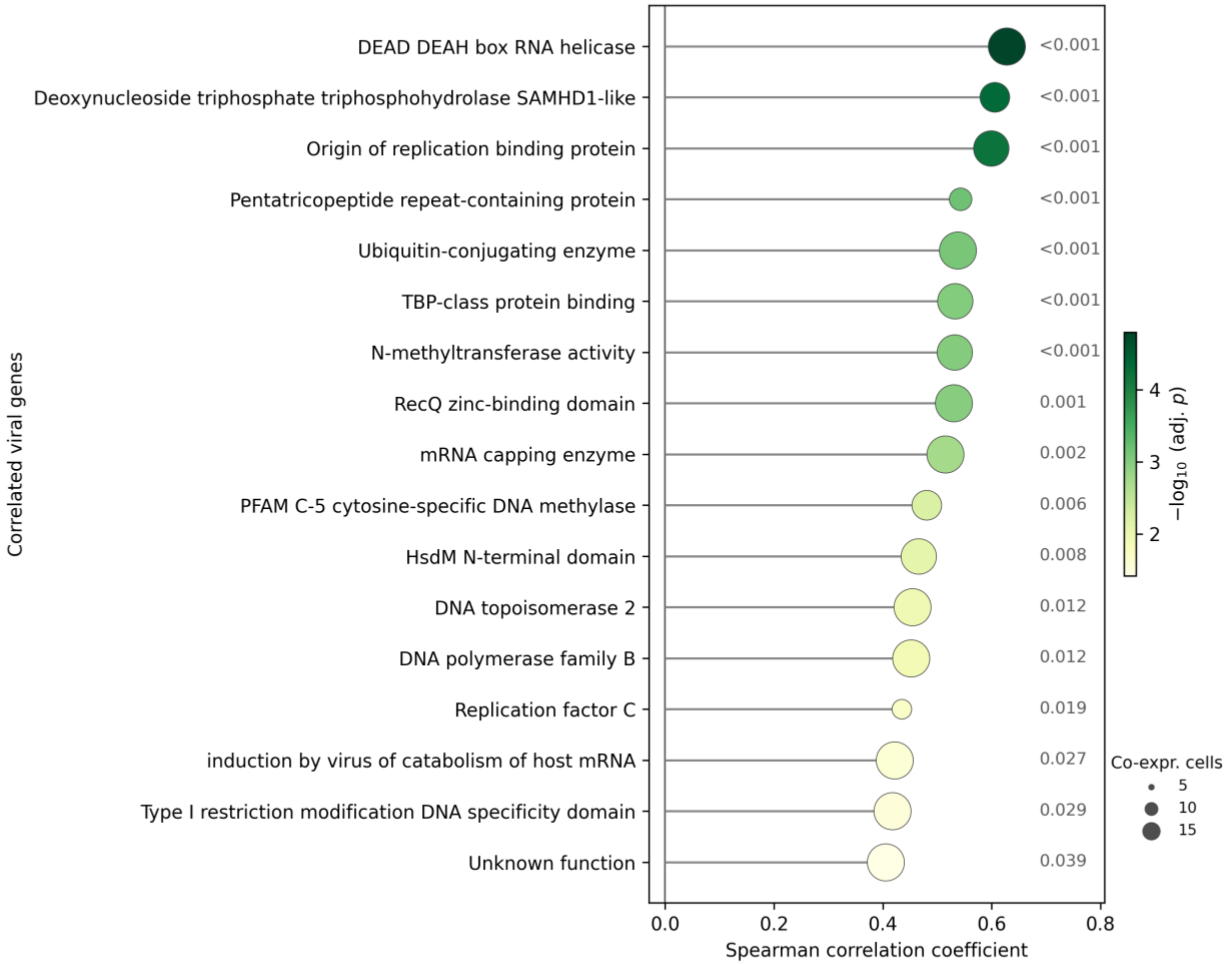


**Fig. S7. Correlation between viral gene expression and viperin expression in single-cell transcriptomes.** Spearman correlation coefficients between the host viperin gene and viral genes across single cells. Genes are ranked by decreasing correlation coefficient. Bubble position on the x-axis indicates the Spearman correlation coefficient (ρ). Bubble color indicates statistical significance, shown as −log10(adjusted *P*), with *P* values adjusted using the Benjamini-Hochberg procedure. Adjusted *P* values are shown to the right of the corresponding points. Only statistically significant correlations are shown.

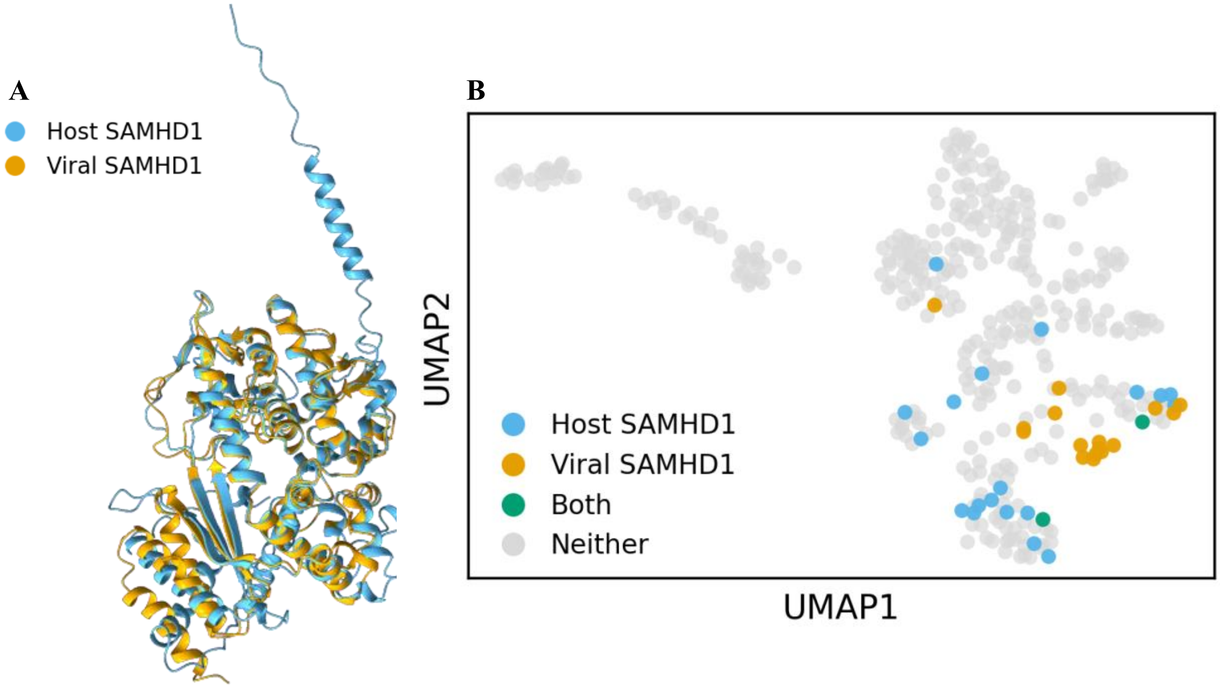


**Fig. S8. Structural similarity and expression of host and viral SAMHD1 in dinoflagellates from Gossenköllesee.** (A) Structural alignment of the putative dinoflagellate host (orange) and viral (blue) SAMHD1 proteins. TM-score: 0.69 (normalized to the host protein) and 0.87 (normalized to the viral protein). (B) UMAP projection of dinoflagellate cells identified by scRNA-seq, showing expression of host and viral SAMHD1 homologs. Cells are colored according to whether they expressed the host SAMHD1 homolog, the viral SAMHD1 homolog, both homologs, or neither.

**Table S1. Summary of samples used for the study**

| Name | Size fraction | Date Collected | Depth | Sequencing method | Used for | Bioproject |
|---|---|---|---|---|---|---|
| APESAF | 0.22µm-40µm | 21/08/24 | 1-8 m composite | Bulk metagenomics | Gene catalog / GVMAGs | PRJNA1483949 |
| AIL3 | 3µm-40µm | 21/08/24 | 1-8 m composite | Bulk metagenomics | Gene catalog / GVMAGs | PRJNA1483949 |
| TPESL | 0.22µm-40µm | 23/08/24 | 7-8 m | Bulk metagenomics | Gene catalog / GVMAGs | PRJNA1483949 |
| TIL3J | 3µm-40µm | 23/08/24 | 7-8 m | Bulk metagenomics | Gene catalog / GVMAGs | PRJNA1483949 |
| GKS0-9 5µm | 5 µm-100 µm | 09/2024 to 08/2025 | 0,4,8 m | Bulk metagenomics | Gene catalog | |
| GKS0-9 0.22µm | >0.22µm | 11/2021 to 11/2022 | 1-8 m composite | Bulk metagenomics | GVMAGs | PRJNA1434480 |
| Alps3 | 3µm-40µm | 21/08/24 | 1-8 m composite | Single-cell RNA-seq | Single-cell RNA-seq | PRJNA1537990 |
| Alps5 | 3µm-40µm | 21/08/24 | 7-8 m | Single-cell RNA-seq | Single-cell RNA-seq | PRJNA1537990 |
| AlpsAmir | 3µm-50µm | 8/23 | 1-8 m composite | Single-cell RNA-seq | Single-cell RNA-seq | PRJNA1537990 |
| scDNA-seq | | | | Single-cell DNA-seq | Single-cell DNA-seq | |

**Data S1. (separate file)**
Relative abundance and taxonomic classification of eukaryotic 18S rRNA sequences detected in the bulk metagenomic samples and used to generate Fig. 2A.

**Data S2. (separate file)**
Taxonomic classification of the 4,389 single amplified genomes (SAGs) recovered by single-cell DNA sequencing, which were classified as eukaryotic.

**Data S3. (separate file)**
High-confidence giant virus-host associations identified by single-cell DNA sequencing and used to generate Fig. 2B.

**Data S4. (separate file)**
Taxonomic assignments of cells recovered by single-cell RNA sequencing based on assembled 18S rRNA transcripts. Cells with a taxonomic completeness score <1 were classified as inconclusive and excluded from downstream analyses.

**Data S5. (separate file)**
High-confidence active giant virus-host associations identified by single-cell RNA sequencing and used to generate Fig. 2C.

**Data S6. (separate file)**
Phylogenetic information for Gossenköllesee metagenome-assembled giant virus genomes (GVMAGs) and reference giant virus genomes used to generate Fig. 3.

**Data S7. (separate file)**
Source data for the Fig. S4 heatmap, including cell barcodes, host taxonomic assignments, viral genomes, and viral transcript counts for individual cells.

**Data S8. (separate file)**
Functional annotations and single-cell expression profiles of genes from the dominant Gossenköllesee giant viruses in Fig. S5, together with sequence and source information for viral and bacterial TIR-nuclease proteins.

**Data S9. (separate file)**
Differential gene expression results comparing infected and uninfected dinoflagellate cells, including the genes shown in Fig. 4, and viral genes correlated with the viperin gene.

**Data S10. (separate file)**
Protein sequences and phylogenetic information used to construct the viperin phylogenetic tree shown in Fig. 5B.
